# Vaccinia Virus Infection Inhibits Skin Dendritic Cell Migration to Draining Lymph Node

**DOI:** 10.1101/2020.02.17.952374

**Authors:** Juliana Bernardi Aggio, Veronika Krmeská, Brian J. Ferguson, Pryscilla Fanini Wowk, Antonio Gigliotti Rothfuchs

**Affiliations:** Department of Microbiology, Tumor and Cell Biology (MTC), Karolinska Institutet, Stockholm, Sweden; Instituto Carlos Chagas, Fundação Oswaldo Cruz (ICC/Fiocruz), Curitiba, Brazil; Department of Pathology, University of Cambridge, Cambridge, UK

**Keywords:** Vaccinia virus, BCG, Dendritic cells, migration, lymph node

## Abstract

Despite the success of Vaccinia virus (VACV) against smallpox there remains a paucity of information on Dendritic cell (DC) responses to the virus, especially on the traffic of DCs and VACV to draining LN (dLN). Herein we studied skin DC migration in response to VACV and compared it to the tuberculosis vaccine *Mycobacterium bovis* Bacille Calmette-Guérin (BCG), another live-attenuated vaccine administered via the skin. In stark contrast to BCG, skin DCs did not relocate to dLN in response to VACV. This happened in spite of virus-induced accumulation of several other innate-immune cell populations in the dLN. UV inactivation of VACV or use of the Modified Vaccinia virus Ankara (MVA) strain promoted DC movement to dLN, indicating that the virus actively interferes with skin DC migration. This active immune suppression by VACV was potent enough to ablate the mobilization of skin DCs in response to BCG, and to reduce the transport of BCG to dLN. Expression of inflammatory mediators associated with BCG-triggered DC migration were absent from virus-injected skin, suggesting that other pathways provoke DC movement in response to replication-deficient VACV. Despite viral suppression of DC migration, VACV was detected in dLN much earlier than BCG, indicating a rapid, alternative route of viral traffic to dLN despite marked blockade of skin DC mobilization from the site of infection.

## Introduction

Dendritic cells (DCs) excel in their capacity to capture, transport and present microbial antigen to prime naïve T cells in secondary lymphoid organs (1). The lymph node (LN) is a major site for such antigen presentation which is often preceded by the relocation of DCs from the site of infection in the periphery to the draining LN (dLN) (2). Despite a large body of data on immunizations with model antigens, DC migration remains incompletely understood during infection with pathogens and live-attenuated bacterial or viral vaccines. Using an infection model in mice and a novel assay to track DC migration *in vivo*, we have previously identified a role for interleukin-1 receptor (IL-1R) signaling in mobilizing a sub-population of skin DCs to dLN in response to *Mycobacterium bovis* Bacille Calmette-Guérin (BCG), the live-attenuated tuberculosis (TB) vaccine (3). In particular, we found the population of migratory CD11b^high^ EpCAM^low^ skin DCs to be engaged in BCG transport from its inoculation site in the skin to dLN and in doing so, to prime mycobacteria-specific CD4+ T cells in the dLN (3).

Like BCG, the smallpox vaccine Vaccinia virus (VACV) is a live-attenuated microorganism administered via the skin. Despite many studies on the immune response to poxviruses and countless investigations on anti-viral T-cell priming, there is a knowledge gap on the initial events that unfold *in vivo* in response to VACV. Due to its large genome and replication-cycle features, VACV is readily used as an expression vector and live-recombinant vaccine for infectious diseases and cancer (4–7). Since BCG efficacy is sub-optimal, there is a standing need to improve TB vaccination. Both improved BCG as well as novel vaccine candidates are considered or have been developed, some of which are currently undergoing clinical trials. In the context of VACV vectors, Modified Vaccinia virus Ankara (MVA) expressing the major *M. tuberculosis* antigen 85A (MVA85A) is an example of a clinically-advanced vaccine candidate (8).

Following inoculation of VACV in the skin, infected cells including LN DCs and macrophages (9) can be detected in the dLN within a few hours (9–12). It is not entirely clear how this rapid relocation of virus from skin to dLN occurs, e.g. if through direct viral access to lymphatic vessels or supported through other mechanisms. Similar observations have been reported after skin infection with Zika virus (12) and Semliki Forest virus (13). In contrast, other studies indicate that VACV is largely restricted to its inoculation site in the skin with limited or null relocation of virus to dLN (14, 15). In this regard, VACV has been shown to interfere with fluid transport in lymphatic vessels and as such to curb its dissemination (16). In addition to data on viral traffic to dLN, there is substantive literature on immunomodulatory molecules produced by VACV and on the overall immunosuppressive properties of VACV infection *in vitro* and in mouse models (17). Using an established toolset for investigating DC responses to mycobacteria, we herein compared local BCG-triggered, inflammatory responses in the skin and skin dLN to that of VACV, especially the ability of VACV to mobilize skin DCs into dLN. We found that unlike the early reaction to BCG, VACV actively inhibits skin DC migration to dLN while retaining the capacity to relocate to dLN in the absence of DC transport.

## Materials and Methods

### Mice

C57BL/6NRj mice were purchased from Janvier Labs (Le Genest-Saint-Isle, France) and used as wild-type (WT) controls. P25 TCRTg RAG-1^−/−^ mice expressing EGFP (18) were kindly provided by Dr. Ronald Germain, NIAID, NIH. All animals were maintained at the Department of Comparative Medicine (KM), Karolinska Institutet. Female mice between 8 and 12 wks old were used. Animals were housed and handled at KM according to the directives and guidelines of the Swedish Board of Agriculture, the Swedish Animal Protection Agency and Karolinska Institutet. Experiments were approved by the Stockholm North Animal Ethics Council.

### Vaccinia virus

Vaccinia virus (VACV) Western Reserve (WR) and deletion mutants ΔA49 (19), ΔB13 (20) and ΔB15 (21) (kindly provided by Prof. Geoffrey Smith, Cambridge University) were expanded on BSC-1 cells. Modified Vaccinia virus Ankara (MVA) was expanded in BHK-21. Viral stocks were purified by saccharose gradient ultracentrifugation. Quantification of plaque-forming units (PFUs) from WR and focus-forming units (FFUs) from MVA was performed as previously described (22) with MVA stocks quantified in chicken embryo fibroblasts cells and WR stocks or WR viral load in lymph nodes (LNs) quantified on BSC-1. In some experiments VACV was inactivated with UV radiation (i-VACV) by placing the virus for 2 minutes in a UV Stratalinker 2400 equipped with 365 nm long-wave UV bulbs (Stratagene, USA). UV-inactivation was confirmed by lack of cytopathic effect on BSC-1 cells infected with i-VACV for up to 3 days (data not shown).

### Mycobacteria

*Mycobacterium bovis* BCG strain Pasteur 1173P2 was expanded in Middlebrook 7H9 broth supplemented with ADC (BD Clinical Sciences) as previously described (23). Quantification of mycobacterial Colony-forming units (CFUs) for bacterial stocks and determination of bacterial load in LNs was performed by culture on 7H11 agar supplemented with OADC (BD Clinical Sciences).

### Inoculation of mice

Animals were inoculated in the hind footpad with 30 μL PBS containing unless otherwise stated, 1×10^6^ CFUs of BCG, 1×10^6^ PFUs of VACV or 1×10^6^ FFUs of MVA. i-VACV was used at an amount equivalent to 1×10^6^ PFUs before UV-inactivation. Control animals received 30 μL of PBS only. For footpad conditioning experiments, animals were injected in the footpad with PBS, VACV or i-VACV 24 hrs before receiving BCG into the same footpad. For studying gene expression in the skin, mice were inoculated in the ear dermis with 5 μL PBS containing the same concentration of mycobacteria or virus as above. Control animals received 5 μL PBS.

Assessment of cell migration from the footpad skin to dLN was done as previously described (3, 24). Briefly, animals previously injected with vaccine or PBS where injected 24 hrs before sacrifice in the same footpad with 30 μL of 0.5 mM 5-and 6-carboxyfluorescein diacetate succinimidyl ester (CFSE) (Invitrogen). For assessing migration after 24 hrs, CFSE was injected 2 hrs after vaccine or PBS inoculation. For studying CD4+ antigen-specific T-cell responses, 1×10^5^ LN cells from naïve P25 TCRTg RAG-1^−/−^ EGFP mice were injected i.v. in the tail vein of C57BL/6 recipients in a final volume of 200 μL. Recipients were infected 24 hrs later in the footpad with 30 µL of BCG or virus. Control animals received PBS. In footpad conditioning experiments, recipients received naïve T cells as above and were injected in the footpad 2 hrs later with PBS, VACV or i-VACV. BCG was given the next day and animals sacrificed 6 days after BCG.

### Generation of single-cell suspensions from tissue

Popliteal LNs (pLNs) were aseptically removed, transferred to microcentrifuge tubes containing FACS buffer (5 mM EDTA and 2% FBS in PBS) and gently homogenized using a tissue grinder. The resulting single-cell suspension was counted by Trypan blue exclusion. In certain experiments, an aliquot was taken and subjected to CFU or PFU determinations as described above. LN suspensions were otherwise washed in FACS buffer and stained for flow cytomety. Ears where excised, transferred into Trizol reagent (Sigma) and homogenized in a TissueLyser (Qiagen, USA) for subsequent RNA extraction, below.

### Flow cytometric staining

Single-cell suspensions from pLN were incubated with various combinations of fluorochrome-conjugated rat anti-mouse monoclonal antibodies specific for CD4 (L3T4), CD8 (53-6.7), CD11b (M1/70), CD11c (HL3), MHC-II I-A/I-E (M5/114.15.2), Ly6G (1A8), CD44 (IM7), CD62L (MEL14), CD69 (H1.2F3), Vβ11 (RR3-15) (BD Biosciences), CD326/EpCAM (G8.8), CD103 (2E7) (Biolegend), CD64 (X54-5/7), CD4 (RM4-5) (eBiosciences) for 45 minutes at 4°C in FACS buffer containing 0.5 mg/mL anti-mouse CD16/CD23 (2.4G2) (BD Biosciences). Flow cytometry was performed on an LSR-II with BD FACSDIVA software (BD Biosciences) and the acquired data analyzed on FlowJo software (BD Biosciences).

### Real-time TaqMan PCR

RNA was extracted from ear homogenates and reverse transcribed into cDNA using M-MLV reverse transcriptase (Promega). Real-time PCR was performed on an ABI PRISM 7500 sequence detection system (Applied Biosystems) using commercially-available primer pairs and TaqMan probes for TNF-α, IL-1 α, IL-1β and Glyceraldehyde 3-phosphate dehydrogenase (GAPDH) (ThermoFisher Scientific, USA). The relative expression of the above factors was determined by the 2-△△Ct method in which samples were normalized to GAPDH and expressed as fold change over uninfected controls.

### Statistical analyses

Analyses were performed using GraphPad Prism 6 (GraphPad Software, Inc., USA). One-way Anova with Tukey’s multiple comparison test was used to compare data group means with a cut-off of p <0.05 considered significant.

## Results

### Skin DCs migrate to dLN in response to BCG but not VACV

To begin investigating the early events after VACV infection, we inoculated C57BL/6 wild-type mice in the footpad skin and assessed immune responses in the draining, popliteal LN (pLN). A CFSE fluorochrome-based migration assay was used to track the movement of migratory skin DCs to dLN. (3, 24). We have used this setup in the past to study early responses to BCG, another live-attenuated vaccine and so included BCG as a comparison to VACV. Corroborating our previous results, BCG skin infection triggered migration of skin DCs to dLN. However, in stark contrast to BCG, skin DCs did not relocate to dLN in response to VACV (Fig. 1A-C). The lack of DC movement in response to VACV was independent of viral inoculation dose (Fig. 1A) and the timepoint in which DC migration was investigated (Fig. 1B). The absence of CFSE-labeled (*i.e.* migratory) skin DCs in the dLN of VACV-infected mice was not concurrent with CFSE labeling in other immune-cell populations, suggesting a generalized absence of cells moving to dLN in response to VACV (Fig. 1C).

**FIGURE 1.**
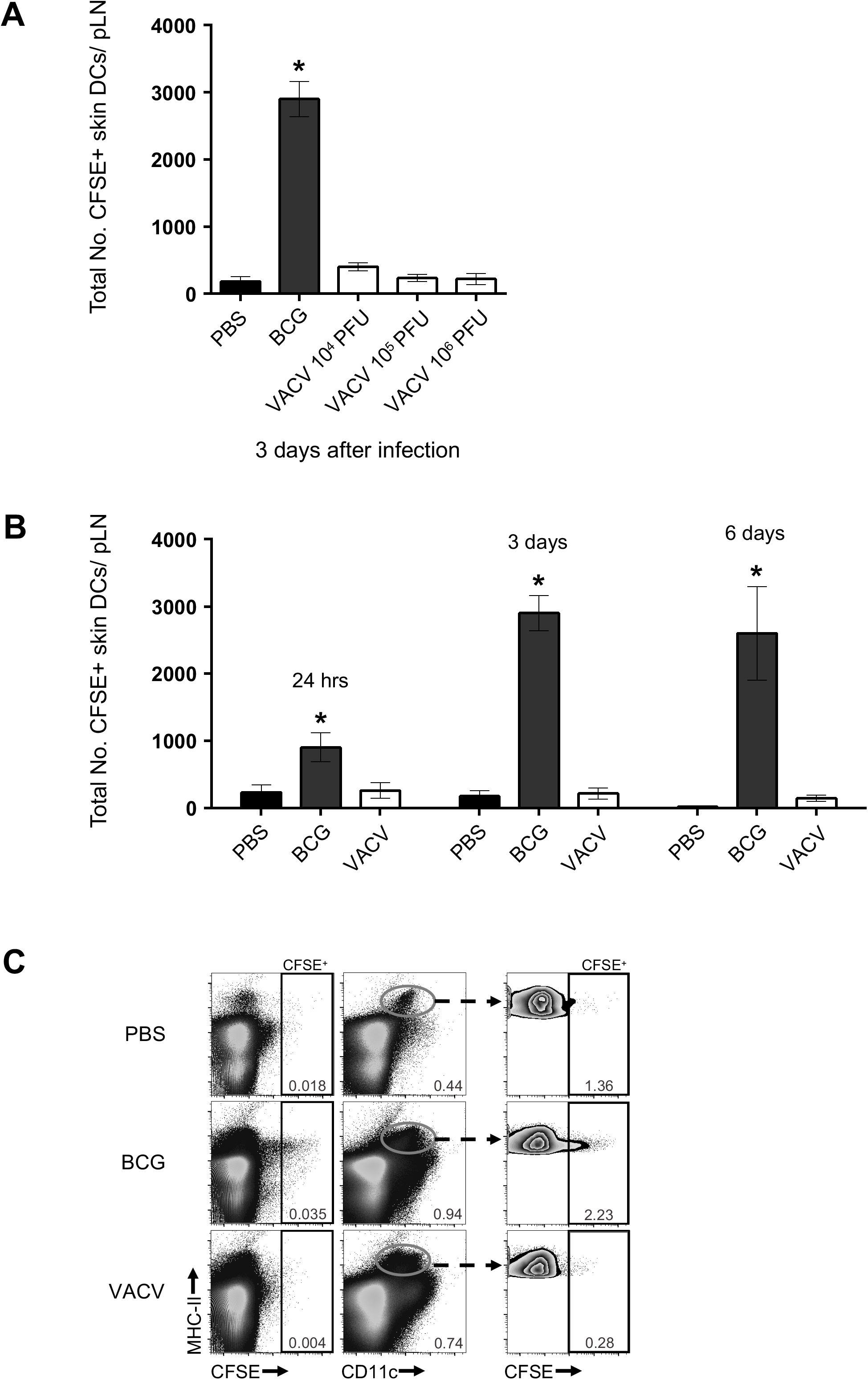
Skin DCs migrate to dLN in response to BCG but not VACV. C57BL/6 mice were inoculated in the footpad skin with PBS, BCG or VACV and subjected to the CFSE migration assay. Single-cell suspensions were generated from the draining, popliteal LN (pLN) and analyzed by flow cytometry. (A) Total number CFSE-labeled skin DCs (MHC-II^high^ CD11c^+/low^) in the pLN 3 days after infection with 10^6^ CFUs of BCG or given doses of VACV. (B) Total number CFSE-labeled skin DCs in the pLN at the given time points after footpad infection with BCG (10^6^ CFUs) or VACV (10^6^ PFUs). (C) Concatenated FACS plots depicting CFSE-labeled cells from pLN 3 days after PBS, BCG (10^6^ CFUs) or VACV (10^6^ PFUs). Representation of CFSE-labeled cells relative to MHC-II (left column). Skin DCs were gated (center column) and CFSE-labeled cells shown (right column). Four to 5 animals per time-point and group used in each experiment. One of two independent experiments shown. Bars indicate standard error of the mean. * denotes statistical difference between PBS and BCG groups.

Although migratory skin DCs did not readily relocate to dLN in response to VACV, virus infection provoked a robust inflammatory response in the dLN, with an increase in LN cell numbers (Fig. 2A) accompanied by the expansion of several innate immune-cell populations and especially CD64^high^ Ly6G^low^ cells (Fig. 2B). The lack of skin DC migration recorded in our CFSE-based migration assay was also in line with a marked decrease in the overall number of migratory skin DCs (MHC-II^high^ CD11c^+/low^ cells) in VACV-infected LN, suggesting that skin DCs may be a particular target of VACV immunosuppression.

**FIGURE 2.**
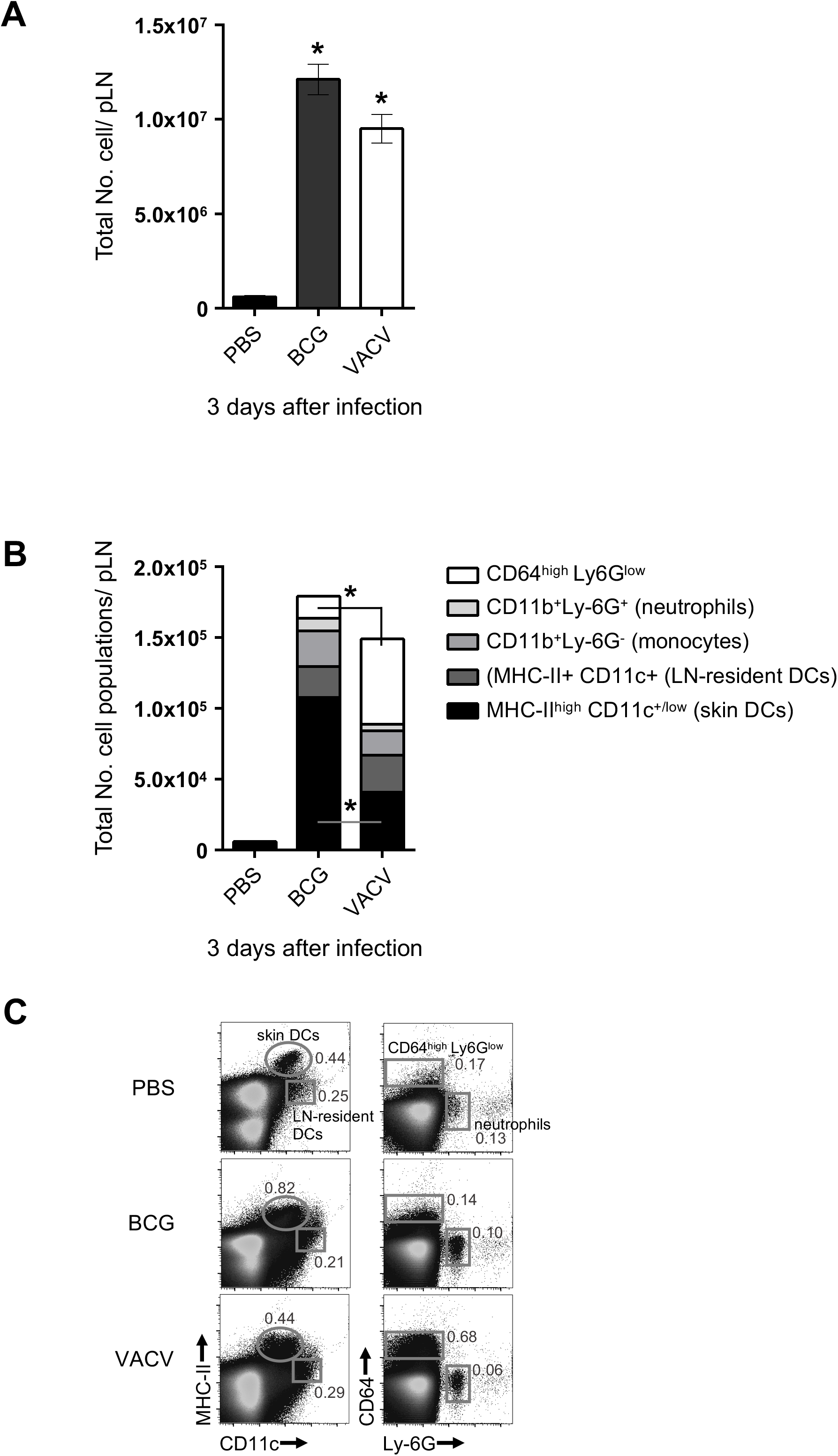
Several innate-immune cell populations expand in the dLN after BCG and VACV infection in the skin. C57BL/6 mice were inoculated in the footpad skin with PBS, BCG (10^6^ CFUs) or VACV (10^6^ PFUs). Three days later, single-cell suspensions were obtained from the pLN and analyzed by flow cytometry. (A) Total cell counts based on trypan blue exclusion. (B). Total number of designated phagocyte populations identified by flow cytometry. Five animals per group were used in each experiment. Bars indicate standard error of the mean. * denotes statistical significance between PBS and infected groups (A) or between CD64^high^ Ly6G^low^ cells and skin DCs, respectively in BCG and VACV groups (B).

### VACV actively inhibits skin DC migration to dLN

Since many of the known immunomodulatory molecules produced by VACV require viral replication, we investigated if the absence of DC migration in response to VACV was coupled to productive infection. We exposed VACV to UV cross-linking at levels sufficient to ablate viral replication but without abolishing viral entry into cells (25). Interestingly, inoculation of UV-inactivated VACV (i-VACV) in the footpad promoted robust skin DC mobilization to dLN (Fig. 3A). Similarly, skin infection with MVA, a highly-attenuated VACV that infects but fails to assemble new virions in mammalian cells (26), also triggered DC migration to dLN (Fig. 3A). Results with i-VACV and MVA thus indicate that live, replication-competent VACV is actively blocking skin DC migration. Moreover, EpCAM^low^ CD11b^high^ DCs were found to be the main DC sub-population contributing to migration in response to i-VACV and MVA (Fig. 3B), an observation similar to BCG (Fig. 3B)(3, 27).

**FIGURE 3.**
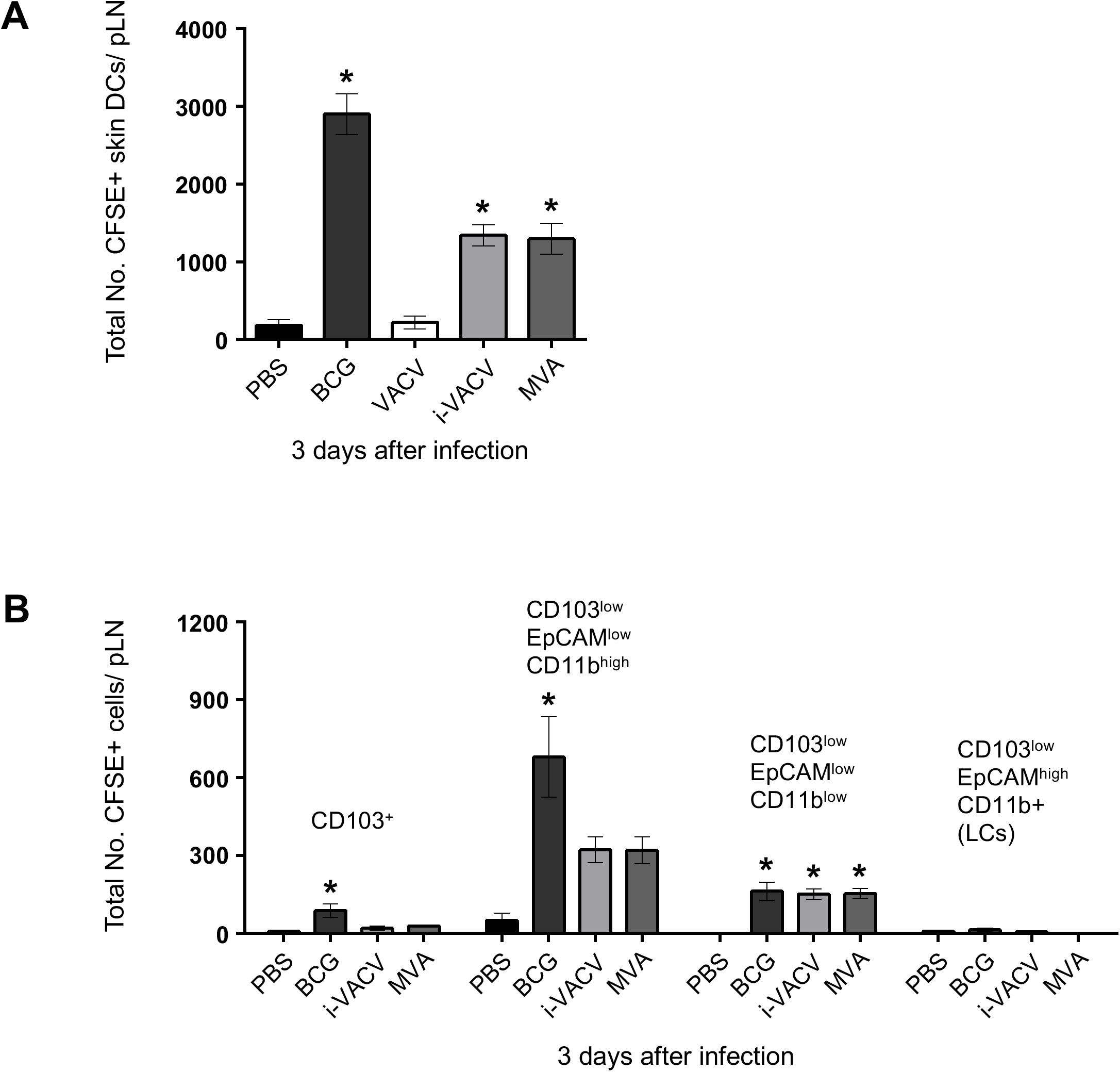
Skin DCs migrate to dLN in response to UV-inactivated VACV and MVA. C57BL/6 mice were inoculated in the footpad skin with PBS, BCG (10^6^ CFUs), VACV (10^6^ PFUs), UV-inactivated VACV (i-VACV, equivalent to 10^6^ PFUs before UV treatment) or MVA (10^6^ FFUs) and subjected to CFSE migration assay as in Fig. 1. pLN were analyzed by flow cytometry 3 days after infection. (A) Total number CFSE-labeled skin DCs shown. (B) CFSE expression within different defined subsets of skin DCs based on previously published gating strategy (3, 24) shown. LC: Langerhans cells. Four to 5 animals per group were used in each experiment. One of two independent experiments shown. Bars indicate standard error of the mean. * denotes statistical significance between PBS- and vaccine-injected groups.

Consistent with a potent suppressive effect on skin DC migration, conditioning the footpad with VACV prior to injecting BCG in the same footpad completely blocked skin DC migration to BCG (Fig. 4A). The absence of skin DCs entering the dLN was also associated with a massive drop in BCG levels in the dLN (Fig. 4B), in line with the fact that these DCs transport BCG to dLN (3). Interestingly, DC migration was not impaired when conditioning was done with i-VACV or MVA (Fig. 4A and data not shown). The number of moving DCs was rather increased (Fig. 4A), although BCG entry itself was slightly lower in the i-VACV-conditioned group compared to PBS-conditioned controls (Fig. 4B). These experiments suggest a carryover of DC migration-dampening properties of VACV infection to a secondary challenge with BCG.

**FIGURE 4.**
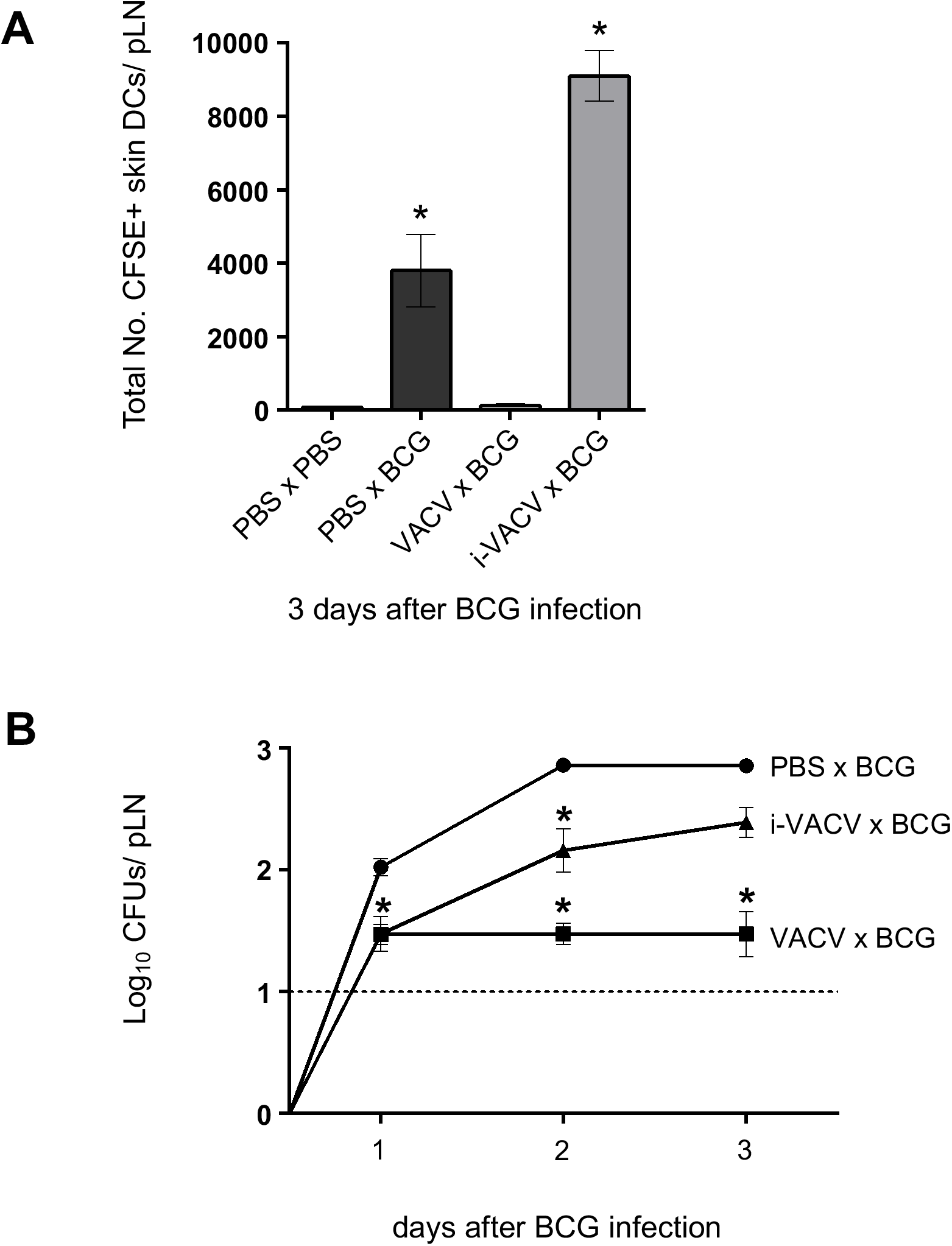
Conditioning the BCG injection site in the skin with VACV mutes the entry of skin DCs and BCG to dLN. C57BL/6 mice were inoculated in the footpad skin with PBS, VACV (10^6^ PFUs) or i-VACV (corresponding to a dose of 10^6^ PFUs before inactivation). Twenty-four hrs later the same footpads were inoculated with BCG (10^6^ CFUs) and the CFSE-based migration assay performed. (A) Total number of CFSE-labeled skin DCs in the pLN 3 days after BCG. (B) Recovery of BCG CFUs from the pLN after conditioning with VACV. Before giving BCG, footpads were inoculated 24 hrs earlier with PBS (PBS x BCG), VACV (VACV x BCG) or i-VACV (i-VACV x BCG). Five animals per group were used in each experiment. One of two independent experiments shown. Bars indicate standard error of the mean. * denotes statistical significance between PBS x PBS controls and vaccine-injected groups (A), or between PBS x BCG and i-VACV x BCG and VACV-BCG groups, respectively.

### Enhanced mRNA expression of inflammatory mediators associated with BCG-triggered DC migration are absent from the skin of VACV-infected mice

Next we compared local changes at the site of infection following inoculation with either vaccine. Enhanced mRNA expression of the pro-inflammatory cytokines TNF-α, IL-1α and IL-1β was clearly detected in the skin 24 hrs after BCG infection (Fig. 5), corroborating our previous observations on a role for IL-1R signaling in regulating DC migration to BCG (3) and (our unpublished data). On the contrary, expression of TNF-α and IL-1 cytokines was absent in response to skin infection with VACV as well as i-VACV and MVA (Fig. 5). Since i-VACV and MVA trigger migration of the same DC subset that also moves in response BCG (Fig. 3B), it is possible that other migration-promoting pathways are in play during i-VACV and MVA-induced migration of EpCAM^low^ CD11b^high^ DCs.

**FIGURE 5.**
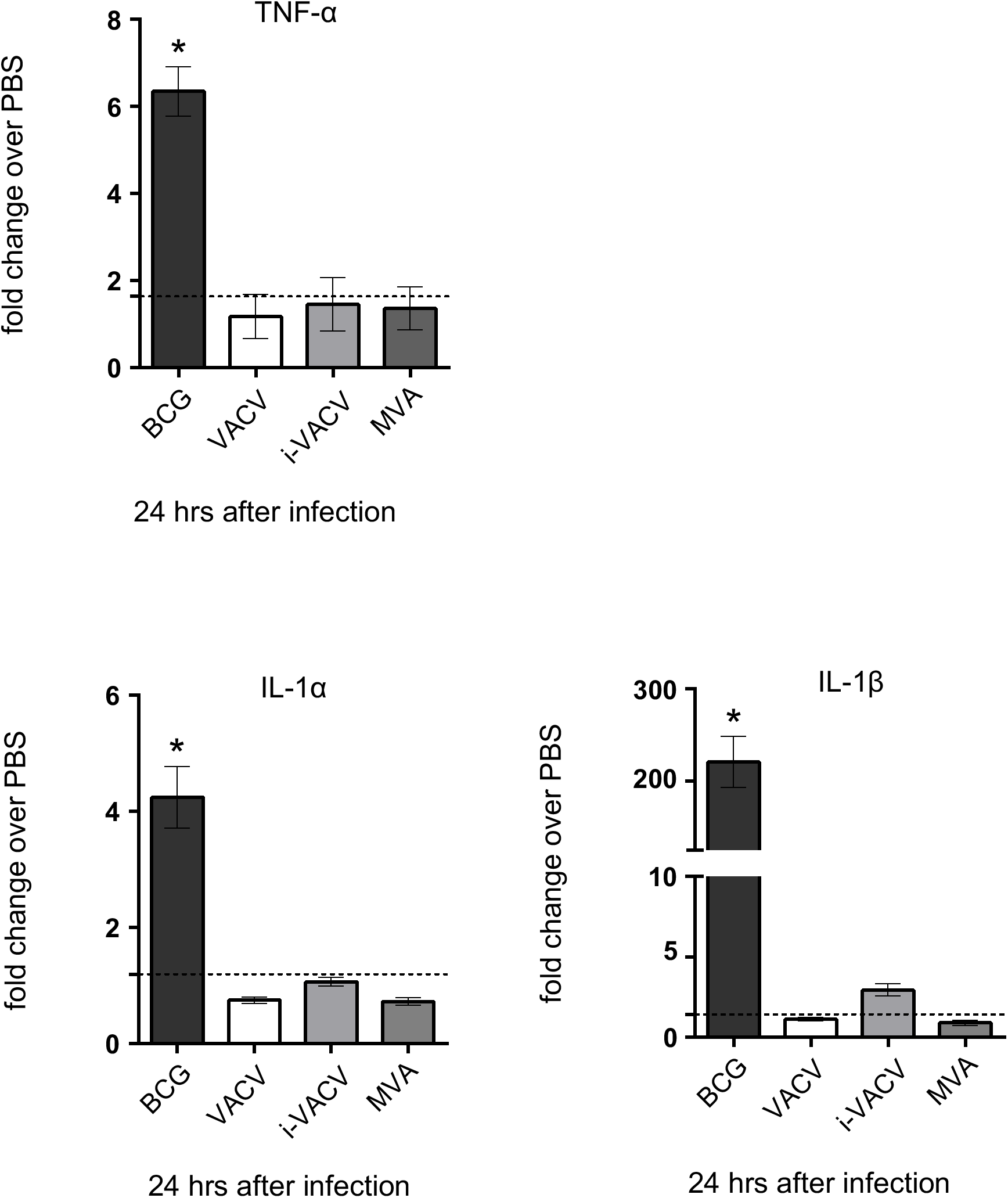
Enhanced mRNA expression of pro-inflammatory mediators associated with BCG-triggered DC migration are absent from virus-infected skin. C57BL/6 mice were inoculated i.d. in the ear with PBS, BCG (10^6^ CFUs), VACV (10^6^ PFUs), i-VACV (equivalent to 10^6^ PFUs) or MVA (10^6^ FFUs). Twenty-four hrs after infection, ears were removed and subjected to RNA extraction and cDNA synthesis. Then mRNA accumulation of TNF-α, IL-1α and IL-1β relative to GAPDH was determined by real-time TaqMan PCR and the fold change of infected animals over PBS controls calculated. Data pooled from 3 independent experiments including 15 to 38 samples per group shown. Dashed line depicts the average relative quantification in the PBS control group for each molecule analyzed. Bars indicate the standard error of the mean. * denotes statistical significance between PBS and BCG groups.

### Early detection of VACV in dLN after infection in the skin

Although DC migration was blocked in response to VACV, the virus was detected in the dLN as early as 10 min after infection in the footpad skin and levels remained steady over time (Fig. 6A). The kinetics of this response was different and notably faster than BCG entry into dLN (Fig 6B), which is known to be reliant on DC transport (3), suggesting an alternative pathway for VACV entry to dLN.

**FIGURE 6.**
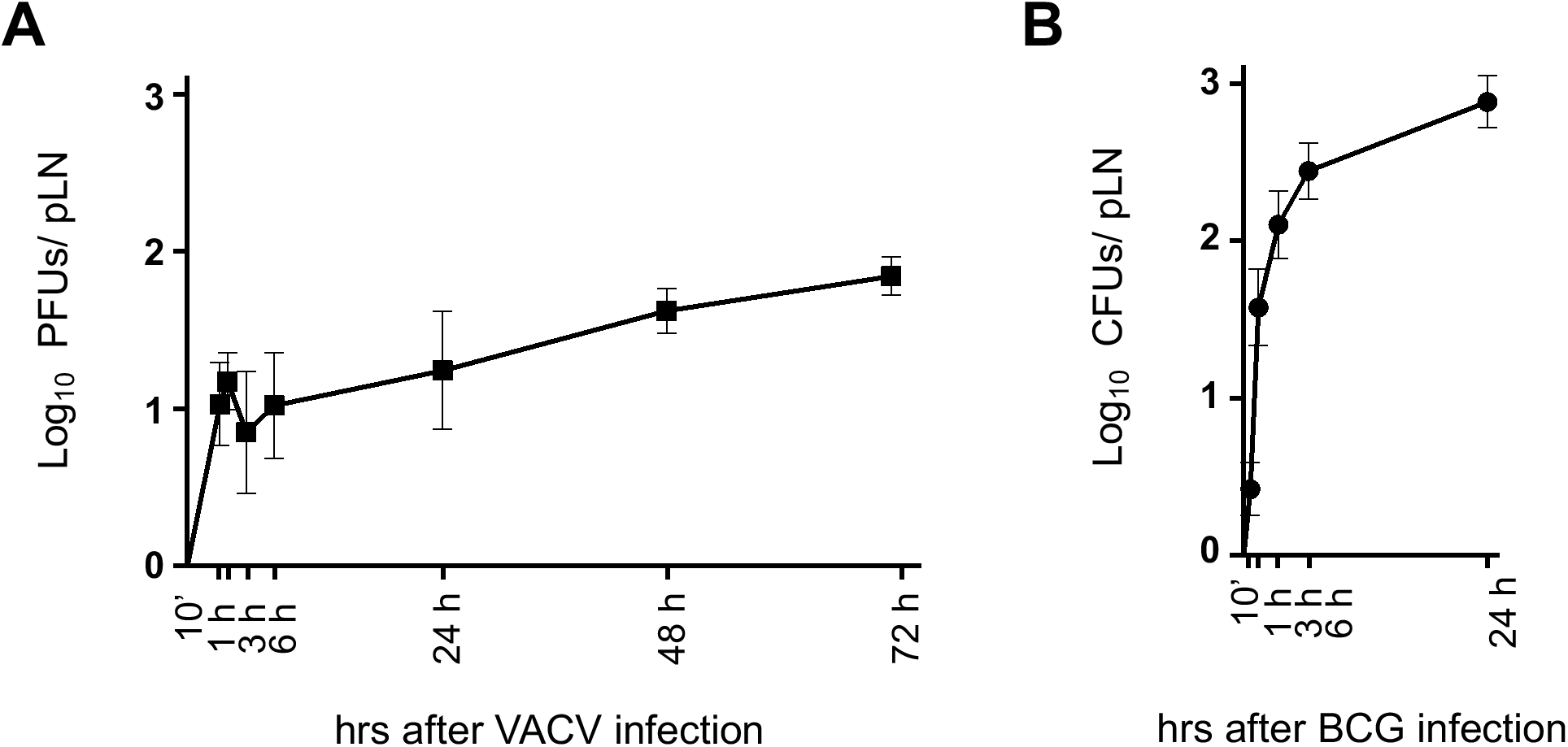
VACV is detected early in dLN after virus infection in the skin. (A and B) C57BL/6 mice were inoculated in the footpad skin with VACV (10^6^ PFUs) or BCG (10^6^ CFUs). Viral (A) and mycobacterial (B) loads in pLN were determined at different timepoints after infection. One of two independent experiment shown. Five animals per time point and group were used in each experiment. Bars indicate standard error of the mean.

## Discussion

VACV is a live-attenuated vaccine that infects a variety of cell types in the skin, including keratinocytes, epidermal and dermal DCs (28). The virus is intriguing in that it packs a diverse immunosuppressive arsenal while still being highly immunogenic. Concurrent with this complexity, the outcome of DC-VACV interactions remains incompletely understood. The fate of the virus and its transport to dLN for antigen presentation is another matter of interest given that VACV is recognized as a vaccine vector and tool for antigen delivery. We show that VACV profoundly inhibits the ability of skin DCs to mobilize to dLN. This inhibition was dependent on viral replication and capable of dampening DC migration to a subsequent challenge with BCG. VACV could nevertheless relocate to dLN in the absence of DC mobilization. Our study supports recent observations that LN conduits transport VACV to dLN for T-cell priming (12). We also add to a large body of data on the immunosuppressive properties of VACV (17) and extend these to include virus-mediated inhibition of DC migration.

VACV infection is known to have many negative effects on DC function. For instance, the virus can inhibit expression of DC costimulatory molecules and cytokines (29–31). Further, splenic DCs isolated from VACV-infected mice have lower MHC-II levels and antigen-presentation capacity (32). We also report lower expression of MHC-II on migratory skin DCs from the dLN of VACV-infected mice (Supplementary Fig. 1A). VACV undergoes abortive replication in DCs (29–31, 33) but VACV and MVA can induce apoptosis in DCs (32). Although we did not assess virus-induced DC death in our studies, the frequency of migratory skin DCs in the dLN of VACV-infected mice was lower than for BCG but similar to that of PBS-injected controls (Supplementary Fig. 1B). This speaks against DC death in the skin, which would lower the pool of migratory DCs in skin and consequently, the frequency of these DCs in the dLN. Results from our migration assay point instead to an impediment of skin DC traffic to dLN during VACV infection.

We also observed muted influx of skin DCs and BCG into dLN if the injection site in the footpad skin had been pre-conditioned with VACV. Thus, the suppressive effect of VACV on DC migration was robust enough to interfere with DC movement triggered by a secondary stimulus (BCG). Interestingly, conditioning with i-VACV doubled the number of migratory skin DCs reaching the dLN without enhancing the entry of BCG. Conditioning with inactivated virus may have reduced skin DC pools available for BCG transport in the next step.

Enhanced expression of TNF-α and IL-1 in the skin was associated with BCG-but not virus-triggered skin DC migration. Migration induced by iVACV and MVA thus highlight an alternative mechanism for EpCAM^low^ CD11b^high^ DCs to move to dLN. Indeed, the contribution of IL-1R signaling in response to BCG is also partial (3) so there must be other factors regulating migration. Similarly, infection with VACV deletion mutants ΔA49, ΔB13 and ΔB15, lacking molecules that inhibit NF-ĸB, Caspase-1 and IL-1 respectively, was not able to provoke DC influx to dLN (Supplementary Fig. 2). During *M. tuberculosis* infection, IFN-α/β has been shown to block IL-1 production from myeloid cells (34). Whether the lack of IL-1 expression in VACV-infected skin or VACV inhibition of skin DC migration is coupled to IFN-α/β remains to be determined. Evaluating cytokine expression and DC migration in IFN-α/βR^−/−^ mice may help clarify this point.

Although VACV did not induce expression of pro-inflammatory cytokines in the skin, it did leash a profound inflammatory infiltrate in the dLN. CD169+ subcapsular sinus (SCS) macrophages are directly exposed to afferent lymph-borne particulates and thus form a strategic line of defense in the dLN against free-flowing viruses, including VACV, preventing their systemic spread (35). Previous studies confirm VACV infection of SCS macrophages (9, 36). In addition, MVA triggers transient inflammasome activation in SCS macrophages that leads to the recruitment of inflammatory cells into the LN (37). In particular, we observed an expansion of CD64^high^ Ly6G^low^ cells in VACV-infected dLN. This population that expanded preferentially in response to VACV remains to be thoroughly characterized. An extensive network of CD64+ macrophages has been reported in the LN paracortex that can scavenge apoptotic cells (38). Early during Listeria infection, populations of inflammatory CD64+ DCs have also been observed in dLN (39).

Migratory skin DCs are tasked with the transport of microbes and their antigens to dLN and constitute as such a central component in our understanding of how adaptive immune responses are initiated in dLN after infection or vaccination in the skin. In our model VACV can reach the dLN without mobilizing skin DCs, whose migration is blocked by the virus. We speculate that skin DC-independent mechanism of virus relocation occurs via direct access of VACV to lymphatic vessels. Previous studies show VACV in dLN within a few hours after injection in the skin (9–12). We report the virus in the pLN even earlier, just a few minutes after inoculation in the footpad. It is unclear how VACV traffics to dLN in the absence of DC transport. The fate of such virus once it arrives in the dLN, including outcome of interactions with LN-resident DCs and ensuing antigen presentation to naïve T cells, remains to be unraveled. Fully unfolding the mechanisms behind VACV blockade of skin DC migration and skin DC-independent relocation of VACV to dLN will profit our understanding of VACV-mediated immune responses and its consequences of T-cell priming.

## Acknowledgements

We thank Susanne Nylén (Karolinska Institutet, Stockholm), for critical reading of this manuscript. We also thank Nuno Rufino de Sousa, Lei Shen, Sören Hartmann and Benjamin Heller Sahlgren (Karolinska Institutet), Marisa Oliveira and Dayana Bozhidarova Hristova (Cambridge University, UK) for technical assistance.

## Disclosure

The authors have no financial conflicts of interest.

---

**SUPPLEMENTARY FIGURE 1.**
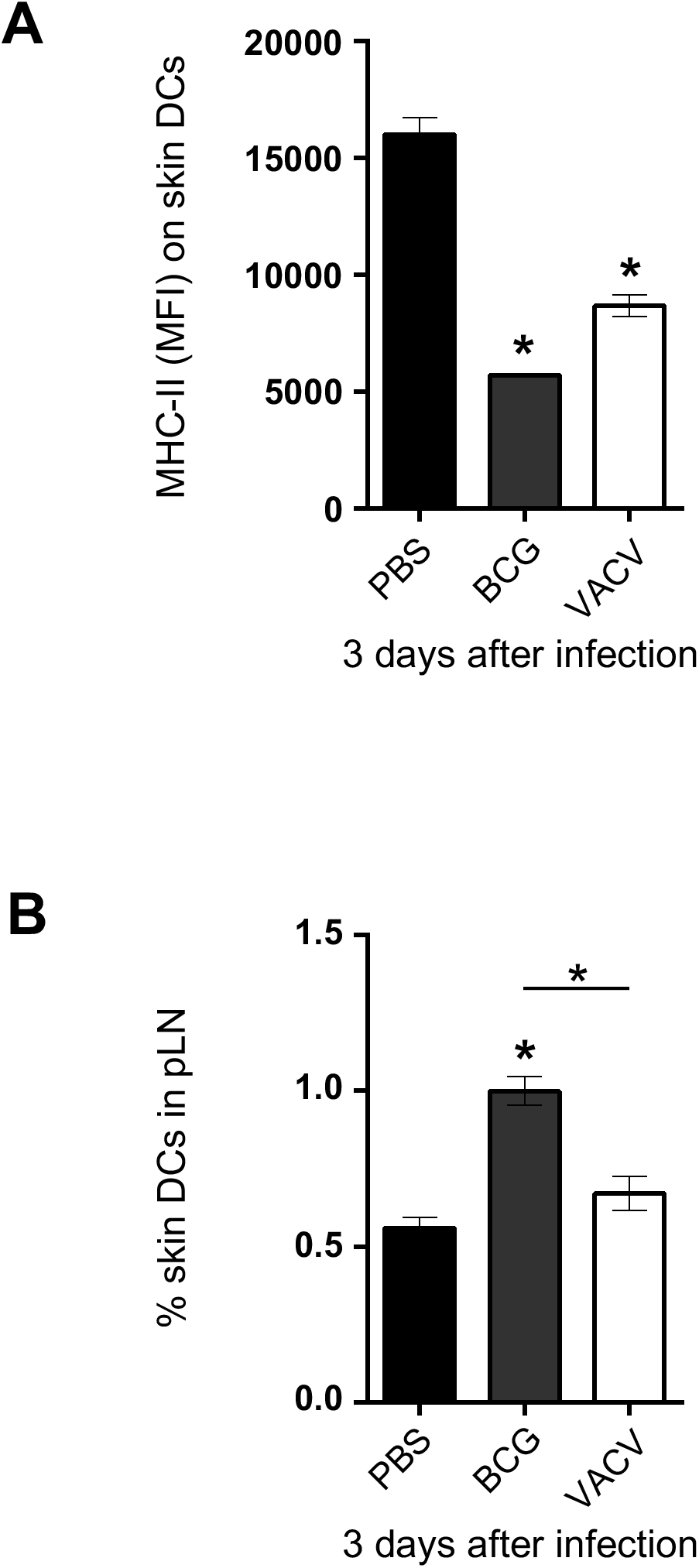
Expression of MHC-II and frequency of skin DCs in dLN of VACV- and BCG-infected mice. (A and B) C57BL/6 mice were inoculated in the footpad skin with PBS, BCG (10^6^ CFUs) or VACV (10^6^ PFUs). Three days later, pLNs were processed and analyzed by flow cytometry. (A) Mean fluorescence intensity (MFI) for MHC-II on skin DCs in pLN. (B) Frequency of skin DCs in pLN. Five animals per group were used in each experiment. One of 3 independent experiment shown. Bars indicate standard error of the mean. * denotes statistically significant difference between PBS and infected groups (A) and PBS and BCG (B).

**SUPPLEMENTARY FIGURE 2.**
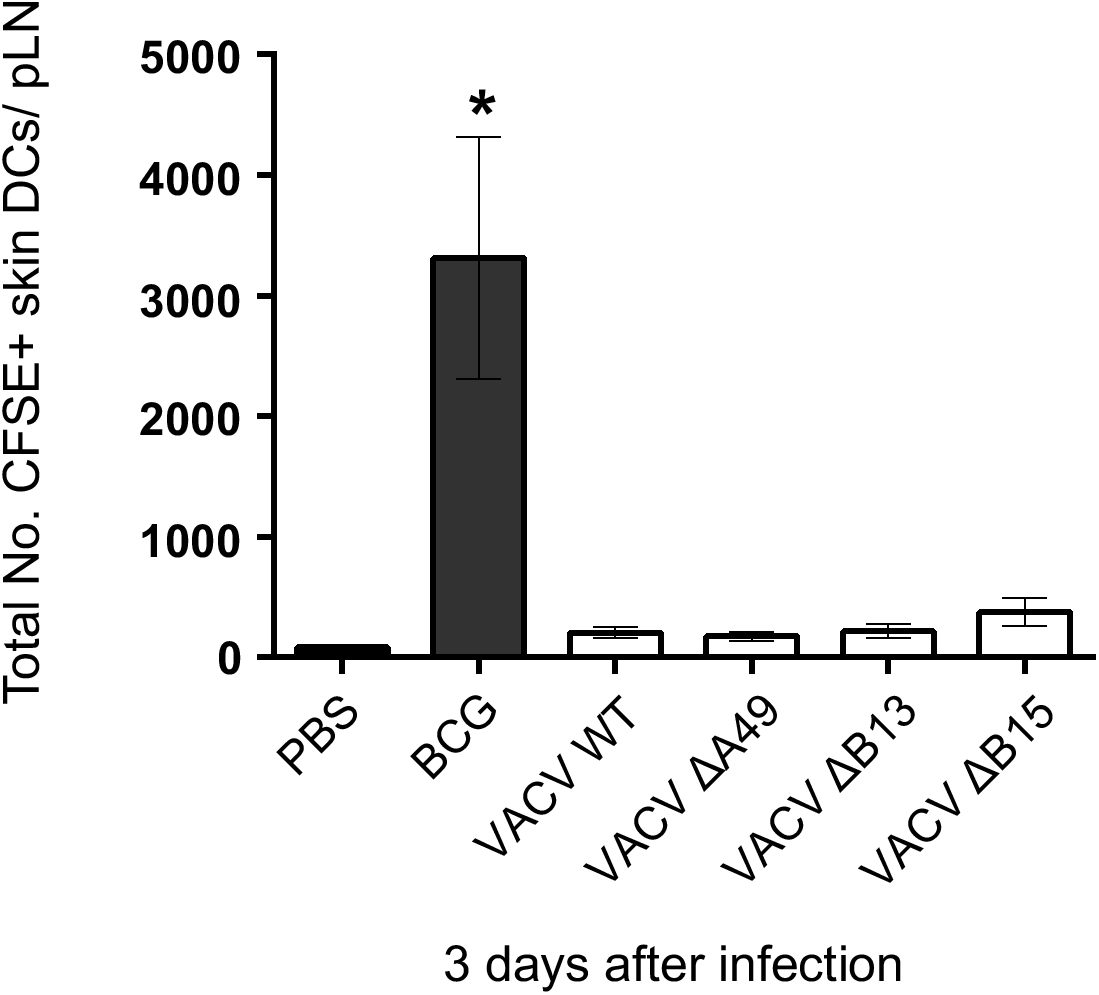
Infection with VACV ΔA49, ΔB13 and ΔB15 does not trigger skin DC migration to dLN. C57BL/6 mice were inoculated in the footpad skin with PBS, BCG (10^6^ CFUs), VACV (10^6^ PFUs) or the VACV deletion mutants ΔA49, ΔB13 and ΔB15 (10^6^ PFUs) and subjected to the CFSE migration assay as in Fig. 1. Total number of CFSE-labeled skin DCs in the pLN 3 days after infection. 5 animals per group were used in each experiment. Bars indicate standard error of the mean. * denotes statistical significance between PBS and BCG groups.

## Notes

Footnotes: This work was supported by the Swedish Research Council (VR) and Karolinska Institutet, Sweden (both to A.G.R), Oswaldo Cruz Foundation (Fiocruz), Brazil (to P.F.W.) and the Foundation for the Coordination for the Improvement of Higher Education Personnel (CAPES), Brazil (to J.B.A. and P.F.W.). The funders had no role in study design, data collection and interpretation or the decision to submit the work for publication.

